# Inactivation of the entire Arabidopsis group II GH3s confers tolerance to salinity and drought

**DOI:** 10.1101/2021.09.24.461718

**Authors:** Rubén Casanova-Sáez, Eduardo Mateo-Bonmatí, Jan Šimura, Aleš Pěnčík, Ondřej Novák, Paul Staswick, Karin Ljung

## Abstract

Indole-3-acetic acid (IAA) controls a plethora of developmental processes. Thus, regulation of their levels is of great relevance for plant performance. Cellular IAA concentration depends on the combined result of its transport, biosynthesis and various redundant pathways to inactivate IAA, including oxidation and conjugation. Group II members of the GRETCHEN HAGEN 3 (GH3) gene family code for acyl acid amido synthetases catalysing the conjugation of IAA to amino acids. However, the high level of functional redundancy among them has hampered thorough analysis of their roles in plant development. In this work, we generated an Arabidopsis *gh3.1,2,3,4,5,6,9,17* (*gh3oct*) mutant to knock-out the group II GH3 pathway. The *gh3oct* plants had an improved root architecture, were more tolerant to osmotic stresses due to locally increased IAA levels and were more drought tolerant. IAA metabolite quantification in *gh3oct* plants suggested the existence of additional GH3-like enzymes in IAA metabolism. Moreover, our data suggested that oxIAA production depends, at least partly, on the GH3 pathway. Targeted stress-hormone analysis further suggested an involvement of ABA in the differential response to salinity of *gh3oct* plants. Taken together, our data provide new insights into the roles of group II GH3s in IAA metabolism and hormone-regulated plant development.

## INTRODUCTION

Regulation of the levels and distribution of the major auxin indole-3-acetic acid (IAA) is of pivotal importance for plant development because it regulates multiple processes, including embryo, seed, root and leaf development, meristem maintenance, shoot branching and responses to environmental stresses (Casanova-Saez & Voss, 2019; Leftley *et al.*, 2021). Spatiotemporal fluctuations of IAA levels in response to external and internal stimuli drive plant developmental responses by triggering signalling cascades that result in modulation of cell division, differentiation and growth (Gallei *et al.*, 2020).

Control of IAA levels and distribution within plant cells and tissues is achieved by intercellular and intracellular transport mechanisms (Band, 2021; Hammes *et al.*, 2021) and spatiotemporally regulated IAA biosynthesis and inactivation (Zhao, 2018; Casanova-Saez *et al.*, 2021). IAA metabolic inactivation primarily involves oxidation processes and the formation of IAA-sugar and IAA-amino acid (IAA-aa) conjugates (Casanova-Saez *et al.*, 2021). IAA oxidation is facilitated by the angiosperm-related DIOXYGENASE FOR AUXIN OXIDATION (DAO) proteins (Zhao *et al.*, 2013; Porco *et al.*, 2016; Zhang *et al.*, 2016; Takehara *et al.*, 2020). The formation of oxIAA is an irreversible IAA inactivation process that operates to remove excess auxin, as inferred from higher oxIAA formation detected at sites of IAA maxima (Pencik *et al.*, 2013). Formation of ester-linked IAA conjugates with glucose (IAA-glc) is a reversible reaction known to regulate IAA levels and homeostasis. Several members of the UDP-glucosyltransferase superfamily have been shown to mediate the conjugation of both IAA and oxIAA to glucose *in vivo* (Mateo-Bonmati *et al.*, 2021).

IAA is also inactivated by conjugation to amino acids via amide bonds, a reaction catalysed by members of the GRETCHEN HAGEN 3 (GH3) family of acyl acid amido synthetases (Staswick *et al.*, 2005). Most of these conjugates, including IAA-leucine, IAA-alanine and IAA-phenylalanine, can be hydrolysed to free IAA, and therefore serve as IAA storage forms (LeClere *et al.*, 2002; Rampey *et al.*, 2004). However, IAA irreversibly conjugates to aspartate (IAA-Asp) and glutamate (IAA-Glu) during catabolism (Ostin *et al.*, 1998; Rampey *et al.*, 2004). IAA-Asp and IAA-Glu catabolites have been found at higher levels compared with reversible IAA-aa conjugates in different plant species (Kowalczyk & Sandberg, 2001; Kojima *et al.*, 2009; Pencik *et al.*, 2009). Several IAA-Asp metabolites, including 6-OH-IAA-Asp and oxIAA-Asp, have been identified in the last few decades (Ostin *et al.*, 1998; Kai *et al.*, 2007). Recently, the IAA oxidase DAO1 was found to additionally operate in the GH3 pathway by mediating the formation of oxIAA-Asp from IAA-Asp in tobacco and Arabidopsis (Müller *et al.*, 2021).

Based on sequence homology and substrate specificity studies, 19 Arabidopsis GH3 proteins have been classified into three functionally diverse groups, of which group II consists of eight GH3 members with the ability to mediate the conjugation of IAA to amino acids, i.e. GH3.1, GH3.2, GH3.3, GH3.4, GH3.5, GH3.6, GH3.9 and GH3.17 (Staswick *et al.*, 2002; Staswick *et al.*, 2005). These group II GH3s are conserved across land plants (Terol *et al.*, 2006).

Even though all Arabidopsis group II GH3s can conjugate IAA to different amino acids, substrate preferences exist among them. For instance, GH3.17 preferentially conjugates IAA to Glu, GH3.3 and GH3.4 preferentially conjugate IAA to Asp, and GH3.2, GH3.5 and GH3.6 show a similar predilection for Asp and Glu (Staswick *et al.*, 2005; Brunoni *et al.*, 2019). A range of amino acid substrate specificities has also been documented for GH3 orthologs in other plant species, e.g. *Physcomitrella patens* (Ludwig-Muller *et al.*, 2009), *Vitis vinifera* (Bottcher *et al.*, 2011) and *Picea abies* (Brunoni *et al.*, 2020). Group II GH3s are also known to be promiscuous regarding the acyl acid substrate. GH3.3, GH3.5 and GH3.6 have been found to modulate jasmonic acid homeostasis by mediating its conjugation to amino acids (Gutierrez *et al.*, 2012). GH3.5 has been further shown to mediate the conjugation of the auxin phenylacetic acid, and the benzoates salicylic acid and benzoic acid (Westfall *et al.*, 2016).

Although specific roles have been reported for individual group II GH3 members (Khan & Stone, 2007; Park *et al.*, 2007; Du *et al.*, 2012; Zheng *et al.*, 2016; Di Mambro *et al.*, 2017; Kirungu *et al.*, 2019), their overlapping expression domains and functions in IAA inactivation have likely masked additional roles of these GH3s in IAA metabolism and plant development. In the present study, we generated a group II *GH3* octuple knockout to bypass the redundancy in GH3-mediated IAA metabolism. The mutant plants showed pleiotropic high-auxin-related phenotypes, such as longer hypocotyls and petioles, as well as a more branched root system with no apparent penalty on primary root growth. Phenotypic, physiological and targeted-hormonomic analyses further indicated that group II GH3s redundantly modulate IAA-dependent salinity tolerance and that they collectively participate in the response to drought. We also obtained evidence supporting a connection between the GH3 pathway and oxIAA production. Finally, our data suggested that additional GH3 or GH3-like genes might operate in IAA conjugation to amino acids.

## MATERIALS AND METHODS

### Plant material and culture conditions

Seeds from the Arabidopsis wild-type accession Col-0 and L*er* and from the *gh3* insertional lines (Table S1) were obtained from the Nottingham Arabidopsis Stock Centre (http://arabidopsis.info) and from the Arabidopsis Biological Resource Center (https://abrc.osu.edu). The presence and position of all insertions were confirmed by PCR amplification using gene-specific primers together with the insertion-specific primers Ds5′-1 (for SGT lines), 3’-dSpm (for SM lines) and LBb1.3 (for SALK lines) (see Tables S1 and S2).

Arabidopsis seeds were surface sterilized with a bleach solution (40% vol/vol commercial bleach in dH_2_O and 0.002% Triton X-100) for 8 min and then washed four times with sterile deionized water. Seeds were stratified for a minimum of 2 days and then sown under sterile conditions on square petri dishes containing half-strength Murashige & Skoog salt mixture (Duchefa, M0221), 1% sucrose, 0.05% MES hydrate (Sigma, M2933) and 0.8% plant agar (Duchefa, P1001) with the pH adjusted to 5.7 witgh potassium hydroxide. Flowering plants were grown in pots containing a 3:1 mixture of organic soil and vermiculite. All plants were grown, unless otherwise stated, under long-day conditions (16 h light, 8 h darkness) at 22 ± 1°C under cool white fluorescent light (150 μmol photons m^−2^ s^−1^). Flowering time was recorded as the number of rosette leaves at bolting under long- and short-day (8 h light, 16 h darkness) conditions.

### Chemical treatments and drought experiments

Salinity and osmotic treatments were performed on half-strength MS plates supplemented with sodium chloride, D-sorbitol or D-mannitol before autoclaving. The inhibitors of YUCCA-mediated IAA biosynthesis yucasin [5–(4–chlorophenyl)-4H-1,2,4–triazole-3–thiol] and PPBo (4-phenoxyphenylboronic acid), kindly provided by Prof. Ken-ichiro Hayashi, were prepared in DMSO stock solutions. IAA treatments were performed on MS plates supplemented with indole-3-acetic acid from a stock solution in absolute ethanol. Yucasin, PPBo and IAA were added to the autoclaved MS media before pouring into plates. To mimic drought conditions *in vitro*, we prepared half-strength MS plates with a reduced water potential as described previously (Verslues *et al.*, 2006). Solidified MS medium was perfused with an overlay solution containing poly(ethylene glycol) (PEG8000; Sigma, P2139). Control plates were perfused with an overlay MS solution lacking PEG8000. For drought treatment on soil-grown plants, stratified seeds were planted in pots containing 40 g of soil mixture. Initially, 1 litre of water was added to each tray. Irrigation was then withheld until both *gh3oct* and wild-type plants displayed dryness symptoms. The weight of the pots was recorded to estimate and compare the water loss in the pots for each genotype.

### Genome-wide identification of T-DNA and transposon insertions

Around 14 μg of nuclear-enriched DNA was purified from 1 g of *gh3oct* seedlings using a previously described protocol (Hanania *et al.*, 2004). Whole-genome sequencing of the sample was performed at BGI Hong Kong using a BGISEQ-500 sequencing platform. 47 million 150-bp-long reads were obtained, reaching a 59X genome depth. Trimmed fastq files were used to map the different insertions using Easymap software (Lup *et al.*, 2021). Raw reads were deposited in the Short Read Archive (SRA) with the code SRX8771538.

### Gene expression analyses

For expression analyses, RNA was isolated using the Total RNA Purification Kit (Norgen). DNA was removed using the RNase-Free DNase I Kit (Norgen). First-strand cDNA synthesis was performed with the iScript cDNA Synthesis Kit (BioRad). *ACTIN2* gene was used as an internal control for relative expression quantification. Four biological replicates (each being a pool of several plants) were analysed in triplicate. qPCR reactions were performed in 10 μl reactions containing 4 μl of LightCycler 480 SYBR Green I Master (Roche), 4 μl of PCR-grade water (Roche), 1 μl of the corresponding primer pair (10 μM each) and 1 μl of the cDNA template. The primers used are listed in Table S2. Quantification of relative gene expression was performed using the comparative C_T_ method (2^−ΔΔCt^) (Schmittgen & Livak, 2008) on a CFX384 Touch Real-Time PCR Detection System (BioRad).

### Quantification of IAA and IAA metabolites

Extraction and purification of the targeted compounds (IAA, oxIAA, IAA-Asp, IAA-Glu, IAA-glc, oxIAA-glc) were performed according to (Novak *et al.*, 2012). Briefly, 10 mg of frozen material per sample was homogenized using a bead mill (27 Hz, 10 min, 4°C; MixerMill, Retsch GmbH, Haan, Germany) and extracted in 1 ml of 50 mM sodium phosphate buffer containing 0.1% sodium diethyldithiocarbamate and a mixture of ^13^C_6_ isotopically labelled internal standards (Olchemim, Olomouc, Czech Republic). After centrifugation (20 000 g, 15 min, 4°C), the supernatant was transferred into new Eppendorf tubes. The pH was then adjusted to 2.5 with 1 M HCl and samples were immediately applied to preconditioned solid-phase extraction columns (Oasis HLB, 30 mg of 1 ml; Waters Inc., Milford, MA, USA). After sample application, each column was rinsed with 2 ml 5% methanol. Compounds of interest were subsequently eluted with 2 ml of 80% methanol. UHPLC-MS/MS analysis was performed according to the method described in (Pencik *et al.*, 2018), using an LC-MS/MS system consisting of a 1290 Infinity Binary LC System coupled to a 6490 Triple Quad LC/MS System with Jet Stream and Dual Ion Funnel technologies (Agilent Technologies, Santa Clara, CA, USA). The quantification was carried out in Agilent MassHunter Workstation Quantitative Analysis software (Agilent Technologies, Santa Clara, CA, USA).

### Quantification of JA, JA-Ile, SA and ABA

Samples were extracted, purified and analysed according to a method described in (Simura *et al.*, 2018). Briefly, around 20 mg of frozen material per sample was homogenized and extracted in 1 mL of ice-cold 50% aqueous acetonitrile (v/v) containing a mixture of [^13^C], [^15^N] or [^2^H] isotopically labelled internal standards using a bead mill (27 Hz, 10 min, 4°C; MixerMill, Retsch GmbH, Haan, Germany) and sonicator (3 min, 4°C; Ultrasonic bath P 310 H, Elma, Germany). After centrifugation (14 000 RPM, 15 min, 4°C), the supernatant was purified as follows. A solid-phase extraction column Oasis HLB (30 mg 1 cc, Waters Inc., Milford, MA, USA) was conditioned with 1 ml of 100% methanol and 1 ml of deionized water (Milli-Q, Merck Millipore, Burlington, MA, USA). After the conditioning steps, each sample was loaded on a SPE column and the flow-through fraction was collected as well as the 1 ml 30% aqueous acetonitrile (v/v) elution fraction. Samples were evaporated to dryness using a SpeedVac SPD111V (Thermo Scientific, Waltham, MA, USA). Prior to LC-MS analysis, samples were dissolved in 40 μL of 30% acetonitrile (v/v) and transferred to insert-equipped vials. Mass spectrometry analysis of targeted compounds was performed using an UHPLC-ESI-MS/MS system comprising a 1290 Infinity Binary LC system coupled to a 6490 Triple Quad LC/MS system with Jet Stream and Dual Ion Funnel technologies (Agilent Technologies, Santa Clara, CA, USA). The quantification was carried out in Agilent MassHunter Workstation Quantitative Analysis software (Agilent Technologies, Santa Clara, CA, USA).

### Phenotypic and statistical analyses

For root and hypocotyl phenotyping, vertically grown plates were imaged using Epson Perfection V600 photo scanners. For petiole length determination, horizontally grown plates were photographed from above. Lengths were measured from scaled images using FIJI software (Schindelin *et al.*, 2012).

Statistically significant differences between mean values from wild type and mutants were analysed by a two-tailed Student’s *t*-test. Multiple comparisons of genotypes, treatments and times were performed by one-way ANOVA followed by Tukey’s post hoc test to a 95% confidence level. Analyses were carried out in GraphPad Prism 6.01.

### Accession numbers

*ACTIN2* (At3g18780), *GH3.1* (At2g14960), *GH3.2* (At4g37390), *GH3.3* (At2g23170), *GH3.4* (At1g59500), *GH3.5* (At4g27260), *GH3.6* (At5g54510), *GH3.9* (At2g47750), *GH3.17* (At1g28130).

## RESULTS

### Arabidopsis group II GH3s are required for vegetative and reproductive development

Group II GH3 enzymes (GH3.1, GH3.2, GH3.3, GH3.4, GH3.5, GH3.6, GH3.9 and GH3.17) are known to redundantly modulate IAA levels by catalysing the formation of IAA-amino acid conjugates, which are inactive IAA forms (Staswick *et al.*, 2005; Chen *et al.*, 2010; Porco *et al.*, 2016). Due to functional redundancy, single *gh3* mutants often show mild or no phenotypes (Staswick *et al.*, 2005). To bypass such redundancy, we generated a *gh3.1,2,3,4,5,6,9,17* octuple mutant (hereafter referred to as *gh3oct*) by crossing the individual insertional lines (Fig. S1a, Table S1). To verify that *gh3oct* was an octuple knock-out, we performed RT-qPCR analysis. We showed that expression of the full-length *GH3* transcript was completely abolished by the insertions in the *gh3oct* mutant, with just marginal expression of *GH3.9* in reproductive tissues (Fig. S1b). Because the average number of T-DNAs in insertional lines has been estimated to be 2.1 (Wilson-Sanchez *et al.*, 2014), the *gh3oct* mutant genome likely harboured several additional insertions that could have jeopardized our interpretations on the GH3-related functions. To rule out this possibility, we sequenced the *gh3oct* genome and followed a tagged-sequence strategy to map the positions of the insertions. We confirmed the presence of eight insertions within the corresponding *GH3* coding sequences and found two extra insertions at intergenic regions (Fig. S2). Due to their position, the latter were considered unlikely to contribute to the *gh3oct* phenotypes.

We then explored the developmental consequences of a group-II *GH3* knock-out. We found that primary roots from *gh3oct* mutant seedlings were comparable in length to those of the wild type, whereas lateral root density was notably increased (Fig. **1a,b,j,k**). Such a root phenotype was observed along the vegetative phase (Fig. **1j,k**). The rosettes from *gh3oct* seedlings showed classical high-auxin phenotypes, such as epinastic cotyledons (Fig. **1a,b**) and elongated hypocotyls (Fig. **1a,b,l**) and petioles (Fig. **1c,d,m**). The *gh3oct* mutant also showed photoperiod-independent early flowering (Fig. **1e**, **S3**). Most of the siliques in the *gh3oct* stems did not elongate properly (Fig. **1i**), likely causing a delayed inflorescence arrest (Ware *et al.*, 2020), and thus making *gh3oct* plants taller (Fig. **1h**). We observed the formation of several aberrant flowers that developed into siliques with unfused valves in the *gh3oct* mutant (Fig. **1f,g**), although this phenotype showed incomplete penetrance. Taken together, our data indicate that the *gh3oct* mutant is very likely an octuple GH3 knock-out at the seedling stage and that functional group II GH3s redundantly control the progress and timing of several developmental processes throughout the plant’s life cycle.

**Figure 1.**
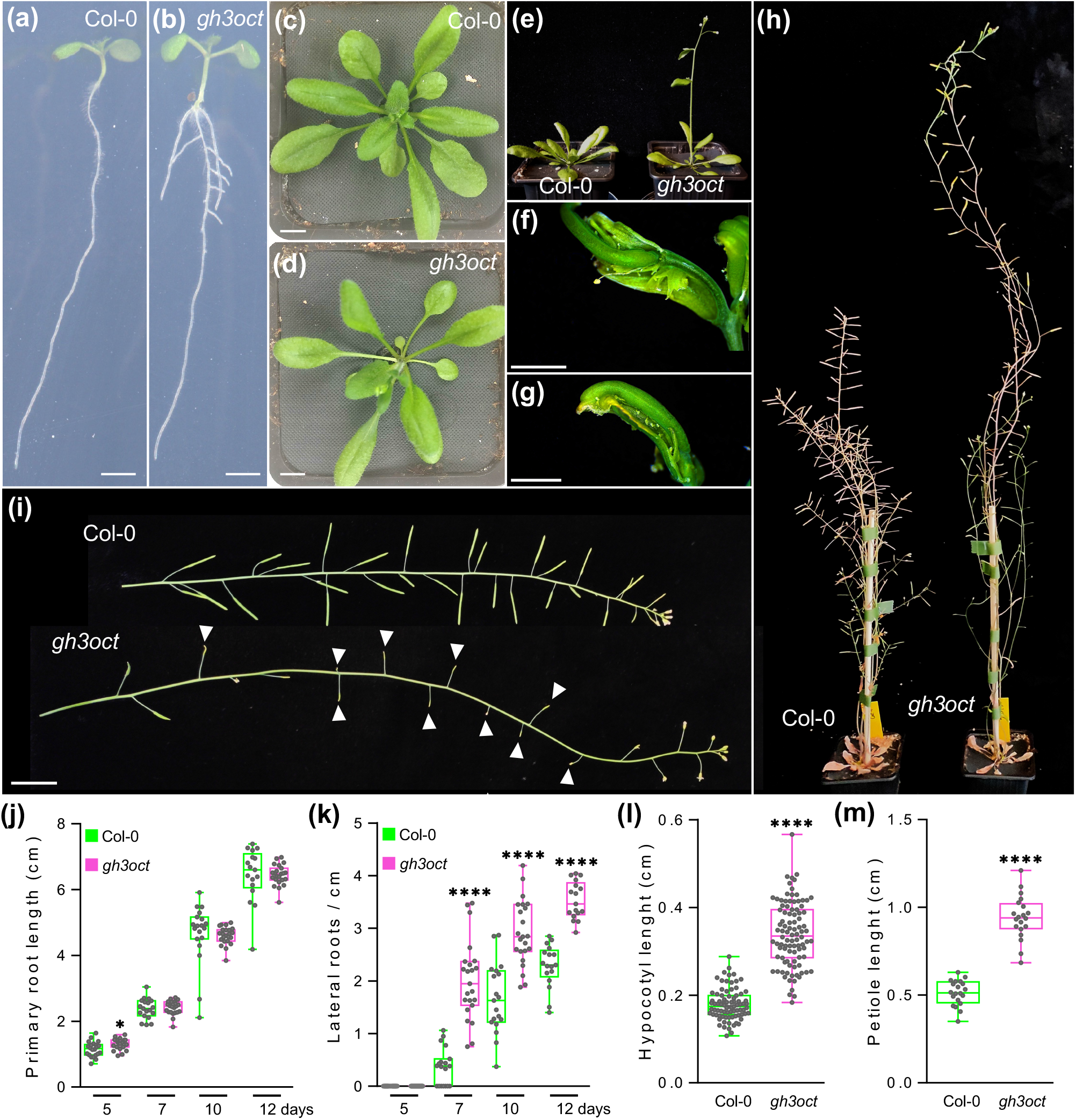
Phenotypes of the *gh3oct* mutant. (a-e) Col-0 and *gh3oct* plants at (a,b) 7, (c,d) 20 and (d) 23 days after stratification (das). (f,g) Aberrant reproductive structures occasionally formed by *gh3oct* mutant from latest inflorescences. (h) Plant height of Col-0 and *gh3oct* flowering plants. (i) Stems from Col-0 and the *gh3oct* mutant. White arrowheads indicate siliques that did not elongate further in the *gh3oct* mutant. (j,k) Primary root length (j) and lateral root density (k) determined at 5, 7, 10 and 12 das. (l) Hypocotyl length determined at 10 das. (m) Petiole length determined at 15 das. Scale bars indicate (a,b) 2 mm, (c,d) 1 cm, (f,g) 2 mm, and (i) 2 cm. Asterisks indicate values statistically different from Col-0 in a two-tailed Student’s *t*-test [\**p*<0.05, \*\*\*\**p*<0.0001; (j,k) *n* ≥ 17, (l) *n* ≥ 83, and (m) *n* = 20].

### IAA metabolism in the absence of functional group II GH3s

Due to the involvement of group II GH3s in IAA metabolism, we explored the concentrations of IAA, the irreversible IAA conjugates IAA-Asp and IAA-Glu, and the IAA inactive forms oxIAA, oxIAA-glc and IAA-glc in shoots and roots from Col-0 and the *gh3oct* mutant. According to the high-auxin phenotypes displayed by *gh3oct* plants (Fig. **1**), IAA contents were found to be increased in *gh3oct* seedling shoots (Fig. **2**). However, total IAA in root tissues was found at wild type levels (Fig. **2**). Despite these modest differences in IAA contents, the roots of *gh3oct* seedlings were hypersensitive to exogenous IAA (Fig. S4). As expected for a total knock-out of the IAA-amido synthetase functions, no detectable IAA-Asp was found in *gh3oct* plants (Fig. **2**). Surprisingly, decreased yet still detectable amounts of IAA-Glu were found in shoots and unchanged IAA-Glu levels were detected in roots from *gh3oct* seedlings (Fig. **2**). The contents of the oxidative catabolites oxIAA and oxIAA-glc were diminished in *gh3oct* shoots and roots (Fig. **2**), whereas no differences in the content of IAA-glc were found (Fig. **2**). Overall, our data suggested that *gh3oct* plants have a reduced ability to inactivate IAA and that IAA inactivation, by means of conjugation to Glu, could still occur in the absence of functional group-II GH3 members. Our data additionally indicated a connection between the production of IAA-aa conjugates and IAA oxidative catabolites.

**Figure 2.**
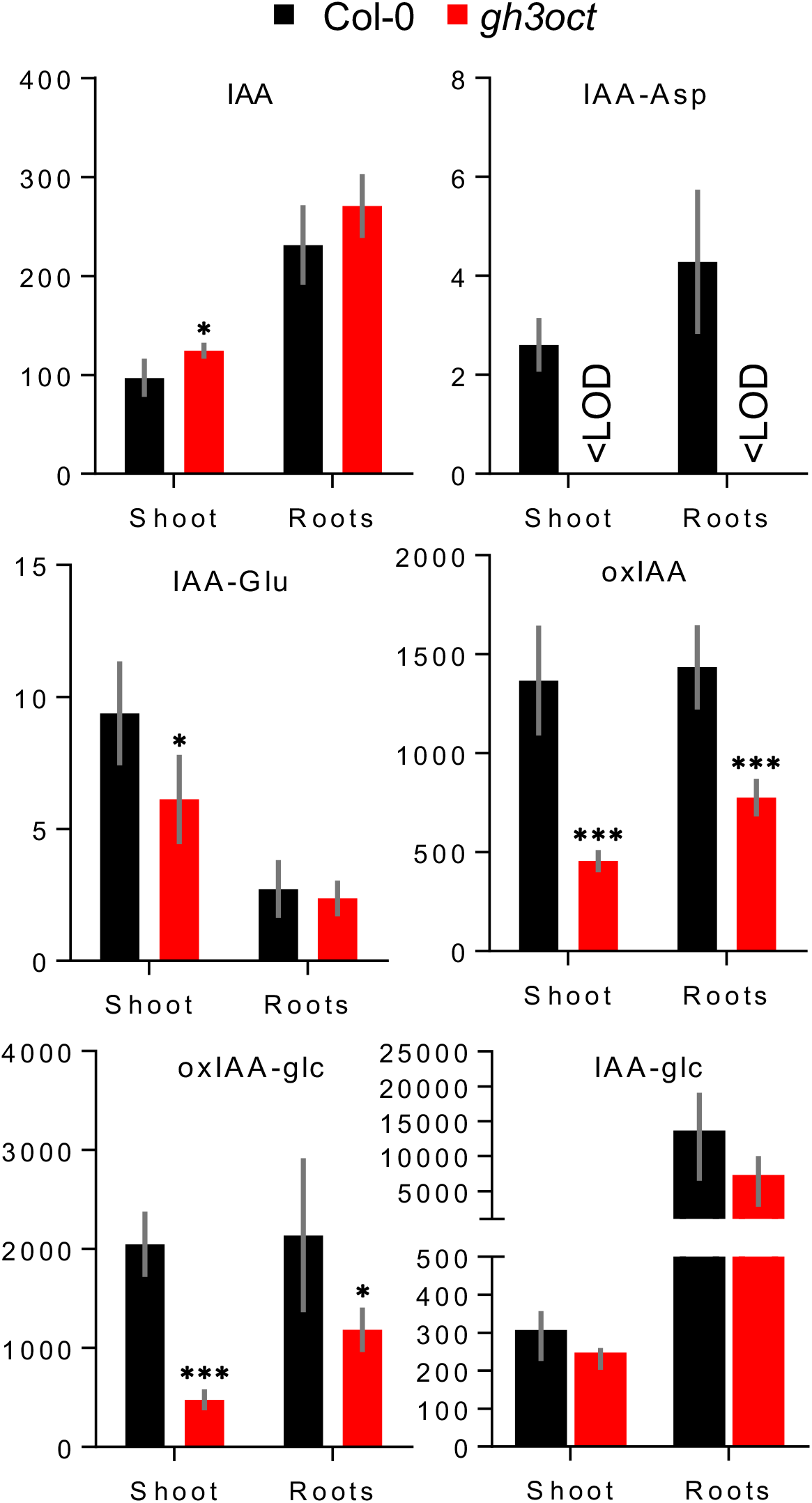
Levels of IAA and IAA metabolites in shoots and roots of the *gh3oct* mutant. Concentration (expressed in picomoles per gram of fresh weight) of indole-3-acetic acid (IAA), indole-3-acetyl-aspartate (IAA-Asp), indole-3-acetyl-glutamate (IAA-Glu), 2-oxindole-3-acetic acid (oxIAA), 1-O-(2-oxindole-3-ylacetyl)-β-d-glucose (oxIAA-glc) and 1-O-indole-3-ylacetyl-β-d-glucose (IAA-glc) in shoots and roots from 7-day-old Col-0 (black bars) and gh3oct (red bars) seedlings. Error bars indicate standard deviation. Asterisks indicate values significantly different from Col-0 in a two-tailed Student’s *t*-test (\**p*<0.05, \*\*\**p*<0.001; *n* = 5).

### *gh3oct* mutant tolerance to salinity and osmotic stress

GH3-mediated IAA metabolism has previously been suggested to modulate abiotic stress responses in plants (Park *et al.*, 2007; Du *et al.*, 2012; Kirungu *et al.*, 2019). Therefore, we focused on the involvement of group II GH3s in the response to salinity. When examining transcriptomic datasets from Arabidopsis plants exposed to NaCl, we found that Arabidopsis group II *GH3s* are responsive to salt treatments (Fig. S5). To corroborate this response under our experimental conditions, we used RT-qPCR to determine the relative expression of each *GH3* gene to NaCl in seedling roots. We found all root-expressed group II *GH3s* to be upregulated upon NaCl application (Fig. **3a**), suggesting that these enzymes might play a role in the response to salinity.

Salinity is well-known to cause general arrest in plant growth (Zhao *et al.*, 2020), and a decreased length of the primary root in salt-stressed Arabidopsis plants is documented (Smolko *et al.*, 2021). Thus, we used the growth of the primary root as an indicator of the tolerance to salinity stress. Compared to Col-0 wild type, *gh3oct* plants were more tolerant to 50 mM NaCl (Fig. **3b,c**). The *gh3oct* mutant also grew better than Col-0 at higher NaCl concentrations (Fig. **3c**). Besides NaCl, we found that the *gh3oct* mutant exhibited higher tolerance to other osmotic stressors, such as sorbitol and mannitol, over the vegetative phase (Fig. S6). Thus, the data suggested that genetic disruption of the group II *GH3s* confers salinity tolerance, likely as part of a general osmotolerant mechanism.

**Figure 3.**
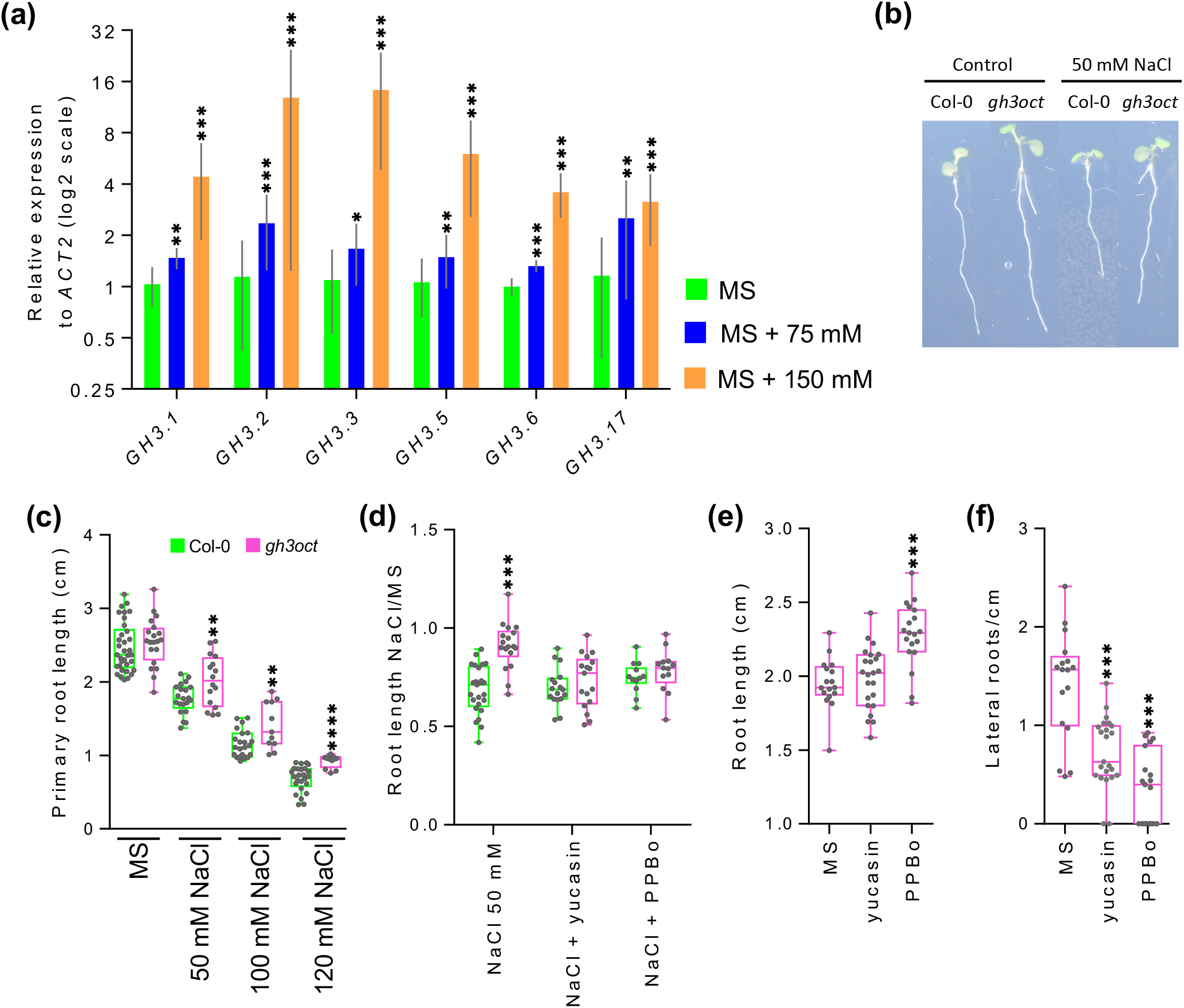
Group II *GH3s* are upregulated upon salt treatment and the *gh3oct* mutant showed IAA-dependent tolerance to salinity. (a) Transcriptional response of group II *GH3* genes from 10-day-old Col-0 roots determined by qRT-PCR 6 hours after transferring to MS media supplemented with 0, 75 or 150 mM NaCl. No expression of *GH3.4* and *GH3.9* was detected in any sample. Error bars indicate standard deviation. (b) Seedlings from 7-day-old Col-0 and *gh3oct* transferred at day 5 to control MS plates or MS plates supplemented with 50 mM NaCl. (c) Primary root length of Col-0 and *gh3oct* seedlings grown for 7 days at different concentrations of NaCl. (d) Root growth ratio of 4-day-old seedlings transferred for another 3 days to media plates containing 50 mM NaCl, 50 mM NaCl + 20 μM of the YUCCA inhibitor yucasin or 50 mM NaCl + 1 μM of the YUCCA inhibitor PPBo (4-phenoxyphenylboronic acid). To calculate the ratio, the length of the root from seedlings grown on NaCl, NaCl + yucasin and NaCl + PPBo plates was divided by the mean root length of seedlings grown on control MS, MS + yucasin and MS + PPBo plates, respectively. (e, f) Root length (e) and lateral root density (f) in 7-day-old *gh3oct* mutant seedlings grown on control MS plates, and on plates supplemented with 20 μM of yucasin or 1 μM of PPBo. Asterisks indicate (a) ΔCT values significantly different from MS (*n* = 4), (c,d) values significantly different from Col-0 (*n* ≥ 11), (e,f) values significantly different from MS (*n* ≥ 17) in a two-tailed Student’s *t*-test (\**p*<0.05; \*\**p*<0.01, \*\*\**p*<0.001, \*\*\*\**p*<0.0001).

### Tolerance to salinity in the *gh3oct* mutant is related to endogenous IAA content

To investigate the relevance of the elevated endogenous IAA content in *gh3oct* plants for the tolerance to salinity, we co-treated *gh3oct* seedlings with both NaCl and IAA synthesis inhibitors. As shown in Fig. **3d**, the higher tolerance to NaCl of the *gh3oct* mutant, in terms of a higher root growth ratio, was abolished by addition of either yucasin or PPBo, which are two inhibitors of YUCCA enzymes that cause a decrease in plant IAA levels by competing with the IAA precursor IPyA (Nishimura *et al.*, 2014; Kakei *et al.*, 2015). To rule out the possibility that the decreased root growth ratio was caused by the effect of the IAA synthesis inhibitors on root growth rather than the tolerance to salinity, we inspected the effects of yucasin and PPBo on the root growth of *gh3oct* seedlings in the absence of NaCl. We found that, at the employed concentrations, none of the inhibitors caused a decrease in the length of the root, whereas both decreased the lateral root density of the *gh3oct* mutant (Fig. **3e,f**).

We next explored the contents of IAA and IAA metabolites and the tolerance to salinity of the *gh3* single mutants. IAA levels were found to be comparable to the wild type in all *gh3* mutant shoots and roots, except for *gh3.1*, which showed decreased IAA levels in roots, and *gh3.2*, which showed increased IAA levels in shoots but decreased levels in roots (Fig. S7). The IAA-Asp concentration was slightly decreased in *gh3.3* shoots and *gh3.6* roots, while a reduced concentration of IAA-Glu was detected in *gh3.1* and *gh3.2* roots (Fig. S7). Levels of oxIAA were decreased in *gh3.2* shoots but were comparable to the wild type in the other *gh3* mutant shoots and roots (Fig. S7). Under our experimental conditions, only *gh3.5* plants showed salt-tolerant root growth (Fig. S8a,b). However, different from the *gh3oct* mutant, the tolerance to salinity of *gh3.5* plants was not suppressed by decreasing endogenous IAA levels (Fig. S8c). Taken together, the results indicated that the tolerance to salinity stress of the *gh3oct* mutant is related to their higher endogenous IAA contents.

### *gh3oct* mutant tolerance to drought stress

Salinity and drought are initially perceived by the plant roots as an osmotic stress. Therefore, it is not uncommon for salt-tolerant genotypes to also exhibit a degree of tolerance to drought stress and *vice versa* (Uddin *et al.*, 2016; Lamers *et al.*, 2020). To test whether *gh3oct* plants better tolerated drought, we first grew Col-0 and *gh3oct* seedlings on media plates with a reduced water potential by using polyethylene glycol (PEG) perfusion (Verslues *et al.*, 2006). The root growth ratio of *gh3oct* plants grown on PEG-perfused plates was higher than that of the wild type (Fig. **4a**), suggesting that the *gh3oct* mutant was more tolerant to drought stress when grown *in vitro*. Because *gh3.5* plants also showed more tolerant growth under salinity, we investigated the growth ratio of *gh3.5* roots in PEG-perfused media. In contrast to the *gh3oct* mutant, *gh3.5* plants were as sensitive as the wild type to PEG-mediated reduction of the water potential (Fig. S8d).

**Figure 4.**
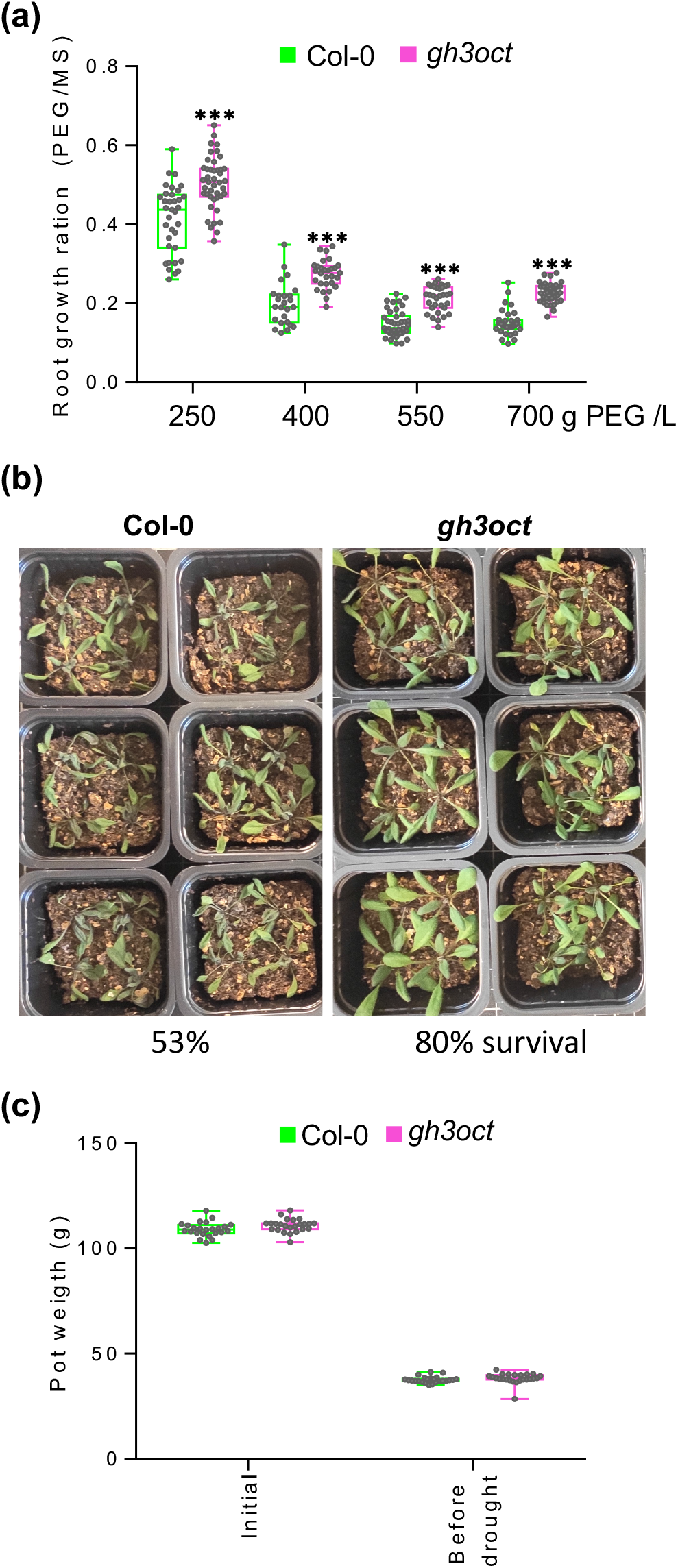
*gh3oct* plants are tolerant to drought. (a) Root growth ratio of Col-0 and *gh3oct* seedlings in media with lowered water potential. 4-day-old seedlings were transferred to plates perfused with a solution of PEG8000 at the indicated concentration and grown for another 6 days. To calculate the ratio, the root length from seedlings in PEG-containing plates was divided by the mean root length of seedlings grown on control plates (perfused with liquid MS). Asterisks indicate statistical significantly different growth ratios from Col-0 in a two-tailed Student’s *t*-test (\*\*\**p*<0.001; *n* ≥ 25). (b) Tolerance of Col-0 and *gh3oct* plants to drought stress on soil. The image corresponds to 3-week-old plants on dry soil. Survival rate after re-watering is indicated at the bottom and represents the mean percentage from 3 independent experiments with 27, 36 and 23 plants per genotype. (c) Soil pot weight in the drought experiment. Pots were prepared with 40 g of soil mixture, watered and weighed at the beginning of the experiment (initial). Pots were re-weighed when the soil was noticeably dry and before plants started showing drought symptoms (before drought). No significant differences were found between Col-0 and *gh3oct* pots in a two-tailed Student’s *t*-test.

Finally, we explored whether the drought tolerance of the *gh3oct* mutant could be reproduced when grown on soil. We planted stratified Col-0 and *gh3oct* seeds in well-watered soil pots and then withheld irrigation until the plants manifested drought symptoms. As shown in Fig. **4b**, *gh3oct* mutant plants showed milder drought symptoms and a higher survival rate after re-watering (80% in *gh3oct* vs. 53% in Col-0). To rule out that pots containing *gh3oct* plants could retain more water, we compared the weight of well-watered to that of dry pots (before drought symptoms appeared) for both genotypes. As shown in Fig. **4c**, we found no differences between Col-0 and *gh3oct* pots. Thus, our results further supported that the *gh3oct* mutant was more tolerant to soil drought.

### Hormonal landscape in *gh3oct* plants under salinity

In addition to IAA, group II GH3 members have been shown to conjugate amino acids to other phytohormones, such as jasmonic acid (JA) and salicylic acid (SA) (Gutierrez *et al.*, 2012; Westfall *et al.*, 2016). Given the involvement of GH3s in the response to salinity, we quantified the endogenous contents of these phytohormones in plants grown in control and NaCl-supplemented media. We found higher IAA levels in *gh3oct* mutant seedlings grown at 0 and 75 mM NaCl, whereas the IAA concentration was comparable to Col-0 at 150 mM NaCl (Fig. **5**). Levels of JA, the JA bioactive form JA-Ile and SA were similar in Col-0 and *gh3oct* seedlings under both control and salinity conditions (Fig. **5**). JA and JA-Ile levels increased in both Col-0 and *gh3oct* plants grown on NaCl-supplemented media (Fig. **5**). Because NaCl stress is known to trigger an increase in levels of abscisic acid (ABA) (Simura *et al.*, 2018), we also quantified levels of this phytohormone. ABA content was higher in *gh3oct* seedlings grown in the control medium, but this difference disappeared in seedlings that germinated under salinity (Fig. **5**).

**Figure 5.**
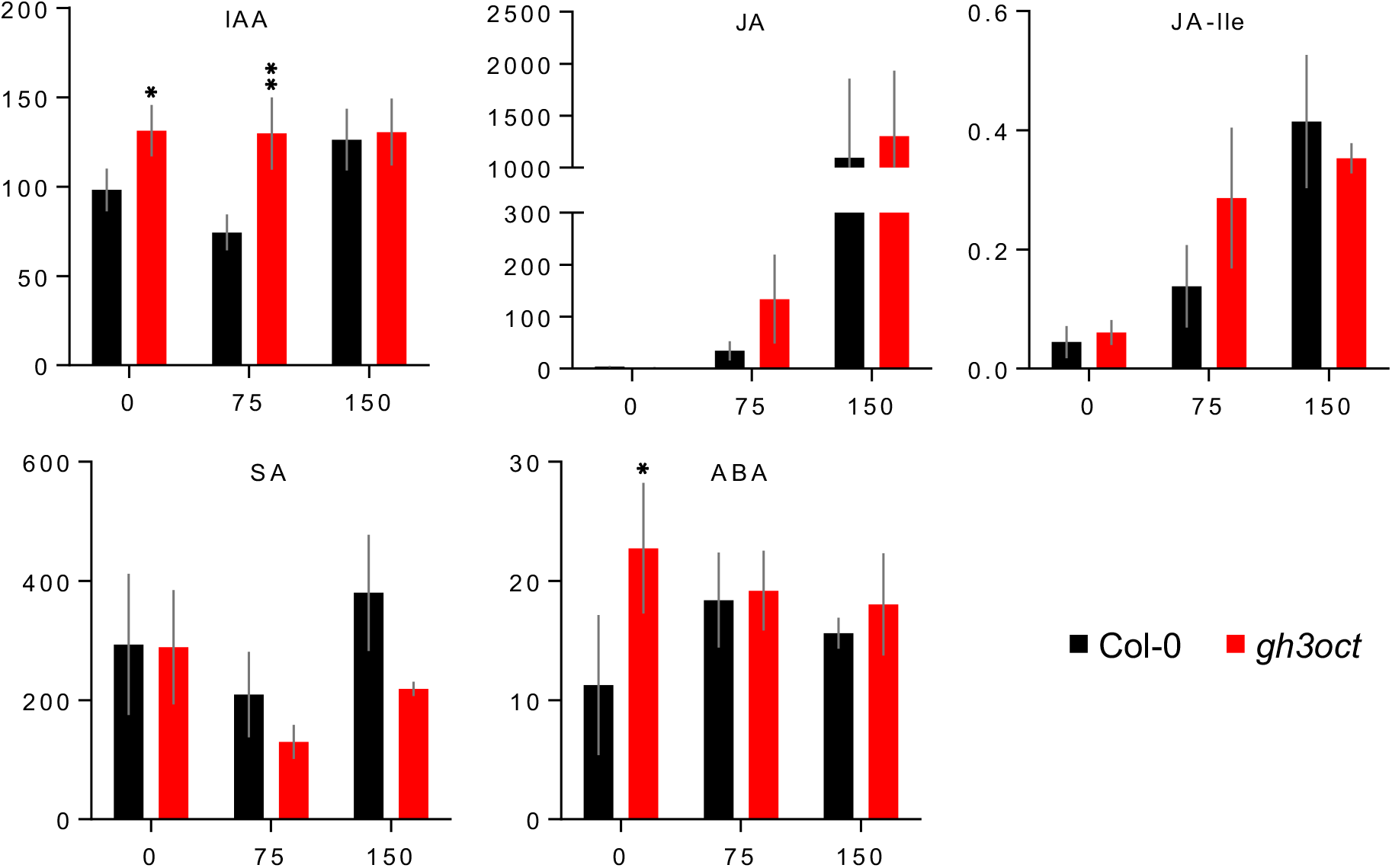
Phytohormone levels from Col-0 and the *gh3oct* mutant grown under different NaCl concentrations. Concentrations, expressed in picomoles per gram of fresh weight, of indole-3-acetic acid (IAA), jasmonic acid (JA), jasmonic acid-isoleucine (JA-Ile), salicylic acid (SA) and abscisic acid (ABA) in Col-0 and *gh3oct* seedlings were determined 7 days after germination on plates containing 0, 75 or 150 mM NaCl. Error bars indicate standard deviation. Asterisks indicate values significantly different from the corresponding values of Col-0 in a two-tailed Student’s *t*-test (\**p*<0.05, \*\**p*<0.01, \*\*\**p*<0.001; *n* = 4). <LOD: below the limit of detection.

Next, we explored changes in endogenous levels of IAA, JA, JA-Ile, SA and ABA in response to salinity. For that, we transferred Col-0 and *gh3oct* seedlings to media plates supplemented with 150 mM NaCl and quantified the phytohormone concentrations at 6 and 24 h upon exposure to NaCl. IAA levels were higher in *gh3oct* seedlings at all inspected times under control conditions, but they became comparable to those of Col-0 in response to 150 mM NaCl (Fig. S9). This result is in accordance with the similar IAA contents in Col-0 and *gh3oct* seedlings grown at 150 mM NaCl concentration (Fig. **5**). Levels of JA and JA-Ile were neither affected by genotype nor by salinity (Fig. S9). Levels of SA were significantly lower in *gh3oct* seedlings after 6 h of exposure to salinity, although this difference vanished after 24 h (Fig. S9). Endogenous levels of ABA prominently increased in both genotypes in response to salinity. However, the ABA content was higher in the *gh3oct* mutant after 24 h of exposure to NaCl (Fig. S9). Because higher ABA levels were also found in *gh3oct* plants grown under control conditions (Fig. **5**) and ABA is known to modulate the response to osmotic stresses, e.g. salinity, by mediating stomatal closure (Hedrich & Shabala, 2018), we hypothesized a higher ability of the *gh3oct* mutant to deal with water loss. However, our observations indicated that the water loss rate in *gh3oct* detached rosettes was indistinguishable from that of the wild type (Fig. S10).

In summary, we found altered levels of IAA and ABA under control conditions and of SA and ABA in response to salinity in *gh3oct* mutant plants.

## DISCUSSION

### IAA metabolism and plant development without functional group II GH3s

Conjugation of IAA to amino acids is an important metabolic pathway regulating levels of this bioactive auxin and is performed by group II GH3 proteins (Staswick *et al.*, 2005). Studies on single *gh3* mutants have demonstrated the relevance of IAA conjugation for plant developmental processes, such as root meristem size control (Di Mambro *et al.*, 2017) and hypocotyl elongation (Zheng *et al.*, 2016). To identify additional roles of the group II GH3s in auxin metabolism and plant development, we generated an octuple group II *GH3* mutant. Our phenotypical analyses of *gh3oct* mutant plants supported redundant involvement of group II GH3s in lateral root, shoot and fruit development, as well as in vegetative-to-reproductive transition. While our data were not sufficient to support a direct causality between IAA metabolism and early flowering or aberrant fruit development in *gh3oct* plants, the observed increased lateral root density and longer hypocotyls and petioles are well-known high-auxin phenotypes (Sugawara *et al.*, 2009; Gao *et al.*, 2020). Our analyses found that the IAA content was only slightly increased in *gh3oct* shoots but at wild-type levels in roots. This suggests that the high-auxin phenotypes displayed by *gh3oct* plants might arise from highly spatiotemporally localized IAA increases, which do not remarkably alter the total IAA content in shoot and root tissues where other mechanisms of IAA control may be more important.

We previously reported a sextuple *GH3* knockout mutant in which 6 group II *GH3s* (GH3.1-6) were disrupted (Porco *et al.*, 2016). No detectable levels of IAA-Asp were found in the *gh3* sextuple mutant, whereas conjugation of IAA to Glu was upregulated. This was likely attributable to the activity of GH3.17, which is highly specific for the formation of IAA-Glu (Staswick *et al.*, 2005; Brunoni *et al.*, 2019). Therefore, one might expect that formation of both IAA-Asp and IAA-Glu would be abolished after knocking out all group II *GH3s*. However, the presence of detectable IAA-Glu content in *gh3oct* seedlings, which displayed null expression of group-II *GH3* genes, strongly suggests that additional GH3 or GH3-like enzymes work in IAA inactivation by means of conjugation to glutamate in Arabidopsis.

DAO proteins have been shown to regulate IAA levels by mediating its metabolic inactivation through oxidation (Zhao *et al.*, 2013; Porco *et al.*, 2016; Zhang *et al.*, 2016). Levels of the oxidative IAA catabolite oxIAA are known to rapidly increase in response to endogenous or exogenous IAA increases (Kubes *et al.*, 2012; Pencik *et al.*, 2013). However, despite showing locally increased IAA levels, as inferred from their multiple high-auxin phenotypes and hypersensitivity to IAA, reduced levels of oxIAA and oxIAA-glc were observed in *gh3oct* plants. DAOs and GH3s have been suggested to establish parallel and redundant pathways for IAA inactivation, with the GH3 pathway playing a compensatory function, as supported by increased and decreased conjugate levels in the DAO1 mutant and overexpressing plants, respectively (Porco *et al.*, 2016; Zhang *et al.*, 2016). This model has recently been challenged by the finding that IAA-Asp is an *in vivo* substrate for DAO1 oxidase, which additionally accounts for the formation of oxIAA-Asp conjugates (Müller *et al.*, 2021). In a context where DAOs function downstream of group II GH3s, the reduced levels of oxIAA and oxIAA-glc in *gh3oct* plants found in the present work suggest that a significant fraction of the plant oxIAA pool derives from oxIAA-aa conjugates.

### Cooperative involvement of group II GH3s in responses to salinity and drought

Salinity and drought are major constraints on plant growth, and therefore represent serious threats to agriculture and forestry. Data from several studies over the past decade support a link between auxin and plant responses to salinity (Wang *et al.*, 2009; Liu *et al.*, 2015; Korver *et al.*, 2018; Fu *et al.*, 2019) and drought stress (Zhang *et al.*, 2009; Zhang *et al.*, 2012; Shi *et al.*, 2014; Jung *et al.*, 2015; Zhang *et al.*, 2020). Spatiotemporal auxin distribution and auxin sensing modulate adaptive growth responses to NaCl stress (Wang *et al.*, 2009; Liu *et al.*, 2015). Recently, Aux/IAA auxin co-receptors were shown to be required for plant tolerance to water deficit, thus suggesting a central role for auxin in integrating drought signals to elaborate genetic and physiological responses (Shani *et al.*, 2017). Although the pathways that mediate auxin-mediated responses to salinity and drought are only just being elucidated, it appears that auxin may mediate tolerance to water deprivation by regulating several adaptive responses leading to stress avoidance. These include (i) adaptive root growth (Leftley *et al.*, 2021), (ii) ROS scavenging (Fu *et al.*, 2019), (iii) regulation of stomatal apertures (Salehin *et al.*, 2019), and (iv) crosstalk with ABA and other hormones involved in the responses to salinity and drought (Sun & Li, 2014; Yu *et al.*, 2020; Salvi *et al.*, 2021).

Our work revealed that *gh3oct* mutant plants showed more tolerant growth under salinity, and that this tolerance was related to endogenous IAA levels. Tolerance to salinity was also higher in *gh3.5* mutant plants, although this effect appeared to be unaffected by endogenous IAA levels. GH3.5 is well-known to mediate the conjugation of other plant hormones, including SA and JA (Gutierrez *et al.*, 2012; Westfall *et al.*, 2016), suggesting that the salt-tolerant growth of *gh3.5* roots may rather be related to the SA or JA pathways. The increased root branching of *gh3oct* plants without penalty for primary root growth, produced a more robust root system, which might contribute to the better performance of *gh3oct* plants under water stress conditions.

In addition to root system architecture, the increased tolerance of *gh3oct* plants to salinity and drought might be driven by increased ABA content. ABA is a stress-related hormone whose levels increase very rapidly upon salt and drought stress sensing and is well-known for its direct involvement in the response to osmotic stresses, primarily by mediating stomatal closure (Hedrich & Shabala, 2018), although other stomata-independent mechanisms have been reported (Thalmann *et al.*, 2016). The increased ABA content but wild-type water loss rate in *gh3oct* plants grown in control conditions indeed rule out a predisposition for *gh3oct* plants to tolerate water deficit based on lower transpiration, but suggests that other ABA-mediated mechanisms might assist *gh3oct* plants to cope with this stress.

## Conclusions

In this work, we present a group-II *GH3* knock-out as a useful genetic tool to research metabolic pathways regulating levels of the main auxin IAA. It has long been accepted that group II GH3 members carry out conjugation of IAA to amino acids. Our metabolic analyses in *gh3oct* plants indicated that additional GH3s or GH3-like proteins might modulate IAA levels by means of conjugation to glutamate. Additionally, our data suggested that the major IAA catabolite oxIAA is, at least in part, produced from the GH3 pathway.

Examination of mutant *gh3oct* plants also revealed that plant tolerance to water deprivation stresses, such as salinity and drought, is modulated by redundant auxin conjugation. Identifying as many components as possible of a plant’s responses to salinity and drought is important for suggesting targets to guide the design of chemical and/or gene-editing approaches directed to enhance stress tolerance in plants. While the mechanisms behind the auxin-related tolerance to salinity and drought remain to be elucidated, the present work supports strategies based on salinity- and drought-inducible disruption of group II GH3s as a profitable tool to engineer plants that better tolerate water stress.

## SUPPORTING INFORMATION

**Figure S1.** The *gh3oct* mutant is an octuple knock-out.

**Figure S2.** The *gh3oct* mutant harbours two extra T-DNA insertions at intergenic regions.

**Figure S3.** The *gh3oct* is a photoperiod-independent early flowering mutant.

**Figure S4.** Root growth in the *gh3oct* mutant is hypersensitive to IAA.

**Figure S5.** Transcriptional response of group-II *GH3* genes to salinity according to published datasets.

**Figure S6.** Root growth in the *gh3oct* mutant is tolerant to different osmolytes.

**Figure S7.** Levels of IAA and IAA metabolites in shoots and roots from group-II *gh3* single mutants.

**Figure S8.** Salinity and drought tolerance of *gh3* single mutants.

**Figure S9.** Levels of IAA and stress-related phytohormones in response to salinity in the *gh3oct* mutant.

**Figure S10.** Col-0 and *gh3oct* plants lose water at the same rate.

**Table S1.** Insertion lines combined to obtain the *gh3oct* mutant.

**Table S2.** Primer sets used in this work.

## ACKNOWLEDGMENTS

Research in the laboratory of Karin Ljung is supported by grants from the Swedish Foundation for Strategic Research (Vinnova), the Knut and Alice Wallenberg Foundation (KAW), the Swedish research councils VR and Formas. E.M.-B. (JCK-1811) held postdoctoral fellowship from Kempestiftelserna. A.P. and O.N. were financially supported by the Ministry of Education, Youth and Sports of the Czech Republic (European Regional Development Fund-Project “Plants as a tool for sustainable global development” No. CZ.02.1.01/0.0/0.0/16_019/0000827). We also acknowledge the Swedish Metabolomics Centre (http://www.swedishmetabolomicscentre.se/) for access to instrumentation.

## AUTHOR CONTRIBUTIONS

E.M.-B., R.C.-S. and K.L. conceived and designed the research; E.M.-B. and R.C.-S. performed most of the experiments; J.Š. and A.P. performed hormone analyses; P.S. made the *gh3oct* mutant; R.C.-S. prepared the manuscript draft; E.M.-B. and R.C.-S. wrote the manuscript with input from all authors. This research was supported by funds to K.L.

