## Supporting information for "Inactivation of the entire Arabidopsis group II GH3s confers tolerance to salinity and drought"

**(a)**

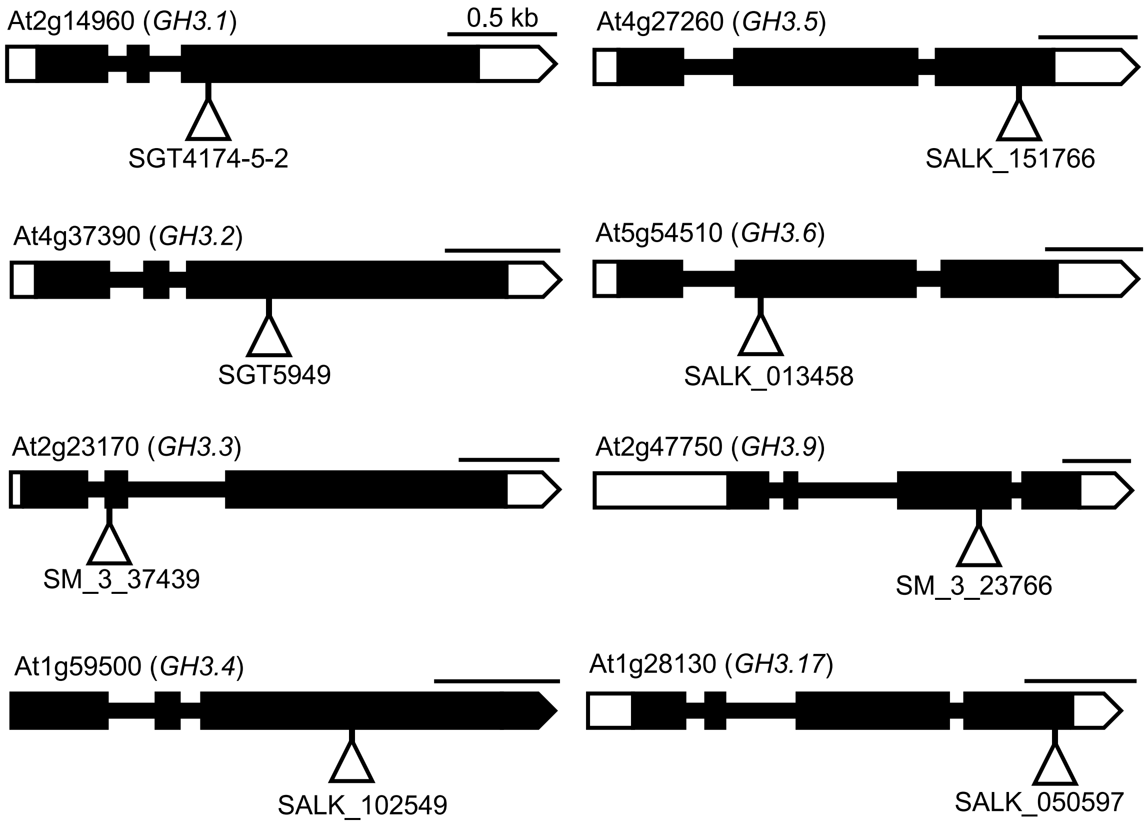


**(b)**

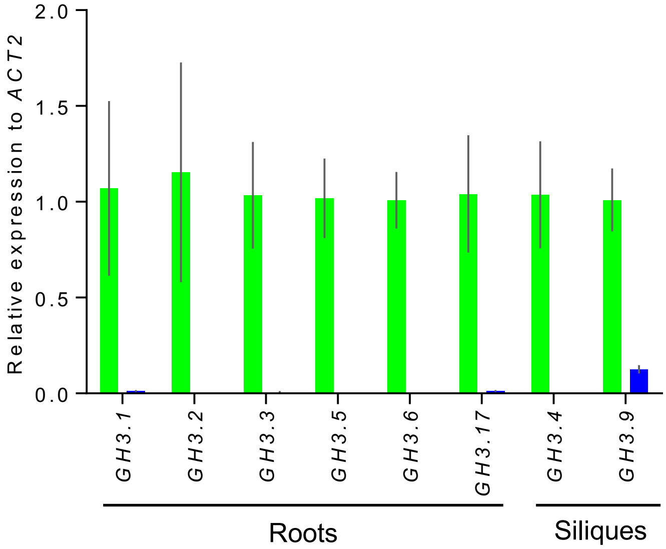


**Figure S1. The *gh3oct* mutant is an octuple knock-out.** (a) Structure of the *GH3.1, GH3.2, GH3.3, GH3.4, GH3.5, GH3.6, GH3.9 and GH3.17* genes with indication of the position of the insertions. Boxes and lines represent exons and introns, respectively. Open and black boxes represent untranslated and translated regions, respectively. Triangles indicate T-DNA/transposon insertions. Scale bars indicate 0.5 kb. (b) Relative expression of *GH3* genes in roots and siliques from 7- and 45-day-old plants, respectively. No expression of *GH3.4* and *GH3.9* was detected in roots. Bars indicate relative expression in Col-0 and the *gh3oct* mutant calculated by using the $2^{-\Delta\Delta CT}$ method. Error bars indicate the interval delimited by $2^{-\Delta\Delta CT \pm SE}$.

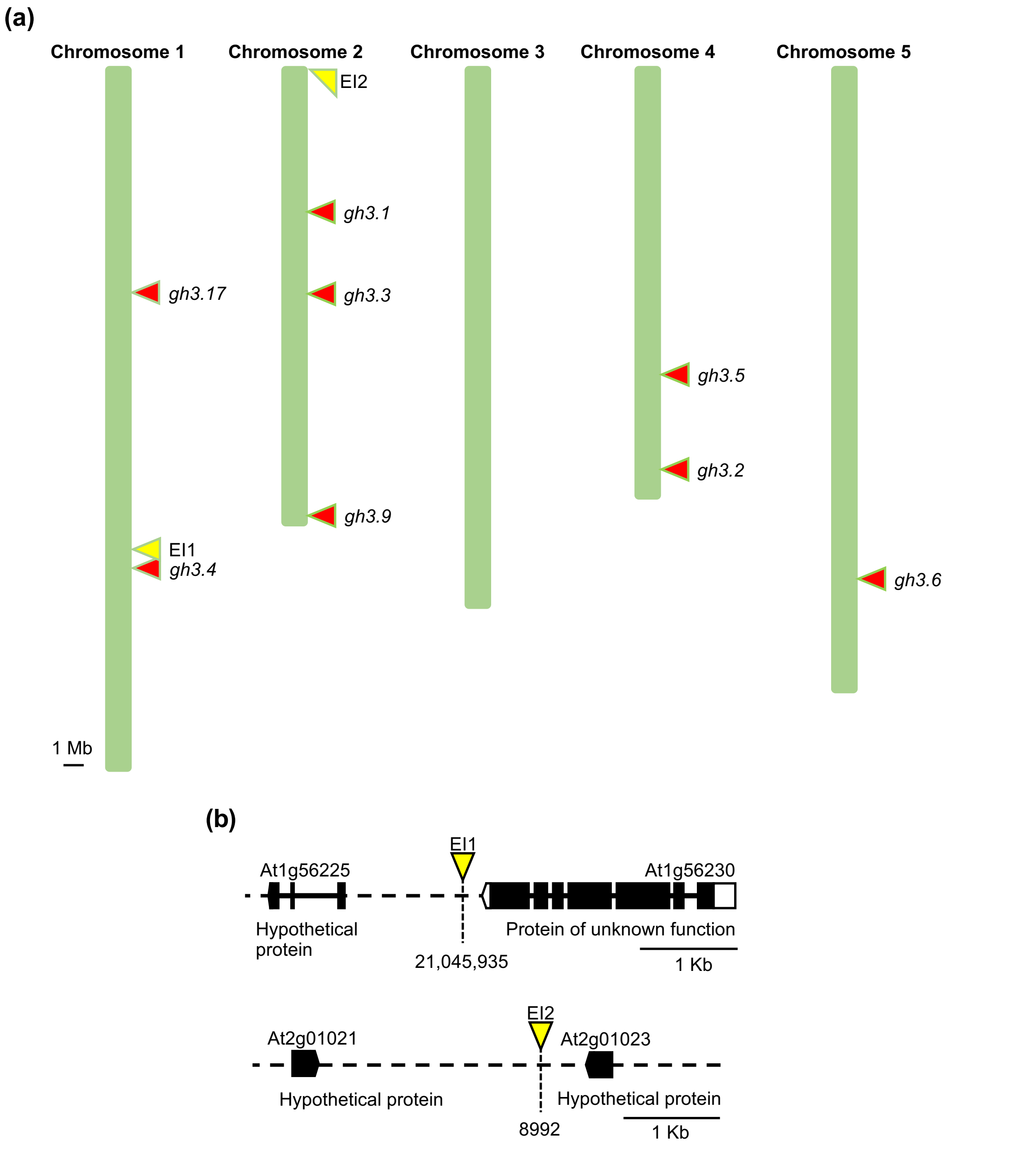


**Figure S2. The *gh3oct* mutant harbours two extra T-DNA insertions at intergenic regions.** (a) Map of the five Arabidopsis chromosomes with indication of the position of the ten insertions found using a tagged-sequence mapping strategy. Insertions disrupting *GH3s* are shown in red. Yellow triangles represent the two extra insertions (EI) found. (b) Detailed locations of the two EI found. Open and black boxes represent untranslated and translated regions, respectively. Scale bars indicate 1 Mb (a) and 1 Kb (b).

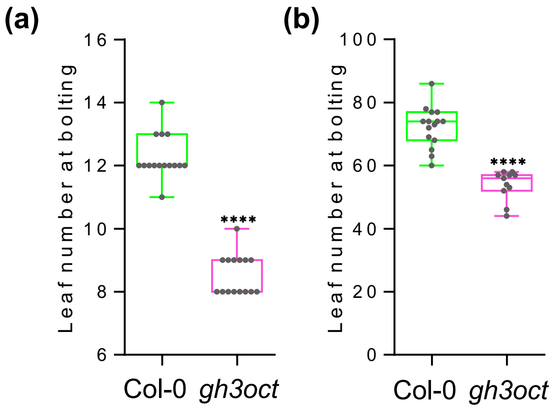


**Figure S3. The *gh3oct* is a photoperiod-independent early flowering mutant.** Flowering time determined as leaf number at bolting in Col-0 and *gh3oct* plants grown under long- (a) and short-day (b) photoperiods. Asterisks indicate values significantly different from those of Col-0 in a two-tailed Student's *t*-test [****$p$<0.0001, $n$ = 15 (a), and $n \geq 11$ (b)].

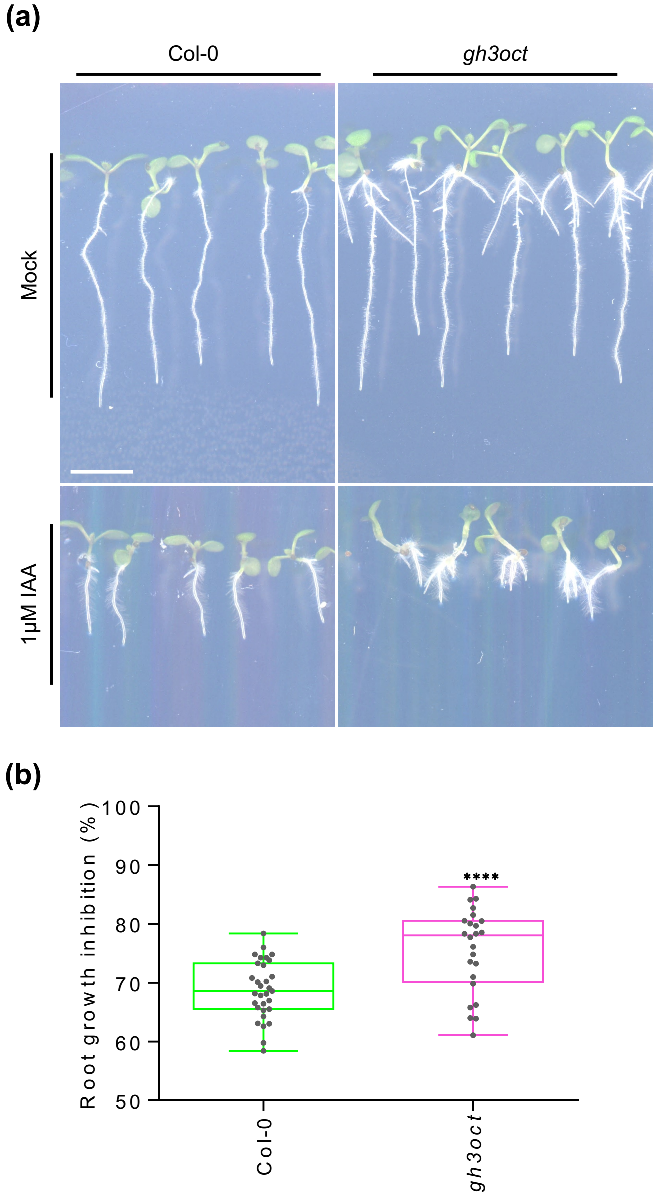


**Figure S4. Root growth in the *gh3oct* mutant is hypersensitive to IAA**. (a) Seedlings from Col-0 and the *gh3oct* mutant on MS plates (top) and MS plates supplemented with 1 μM IAA (bottom). Photographs were taken 7 days after stratification. Scale bar indicates 0.5 cm. (b) Percentage of root growth inhibition by 1 μM IAA in 7-day-old seedlings. Asterisks indicate values significantly different from Col-0 in a two-tailed Student´s *t*-test (****p*<0.001, *n* ≥ 24).

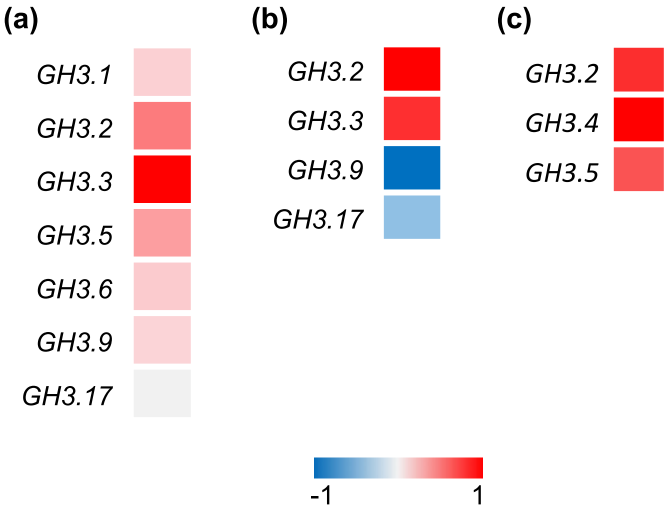


**Figure S5. Transcriptional response of group-II *GH3* genes to salinity according to published datasets.** Data were retrieved from RNA-seq datasets described in (a) Cai *et al.*, 2017 (b) Shen *et al.*, 2014 and (c) Jiang and Deyholos, 2006. Colour scale indicates normalized (a) RPKM and (b,c) $log_2$ fold change values.

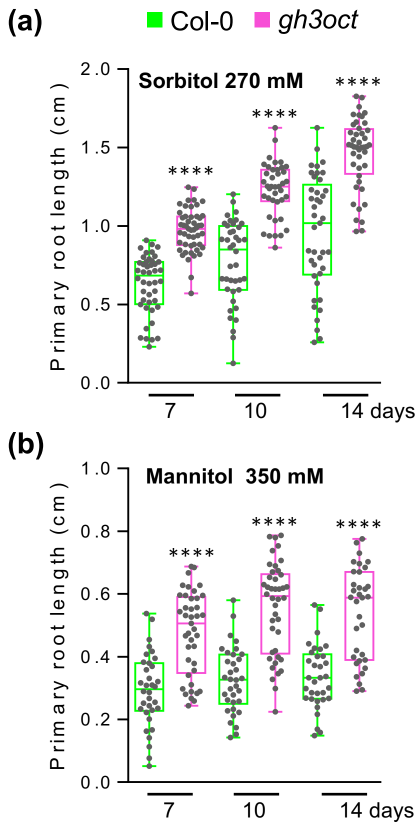


**Figure S6. Root growth in the *gh3oct* mutant is tolerant to different osmolytes.** Primary root length from Col-0 and *gh3oct* seedlings grown on MS plates supplemented with (a) 270 mM sorbitol, and (b) 350 mM mannitol for 7, 10 and 14 days. Asterisks indicate values significantly different from those of Col-0 in a two-tailed Student´s *t*-test (****$p$<0.0001, $n \geq 33$).

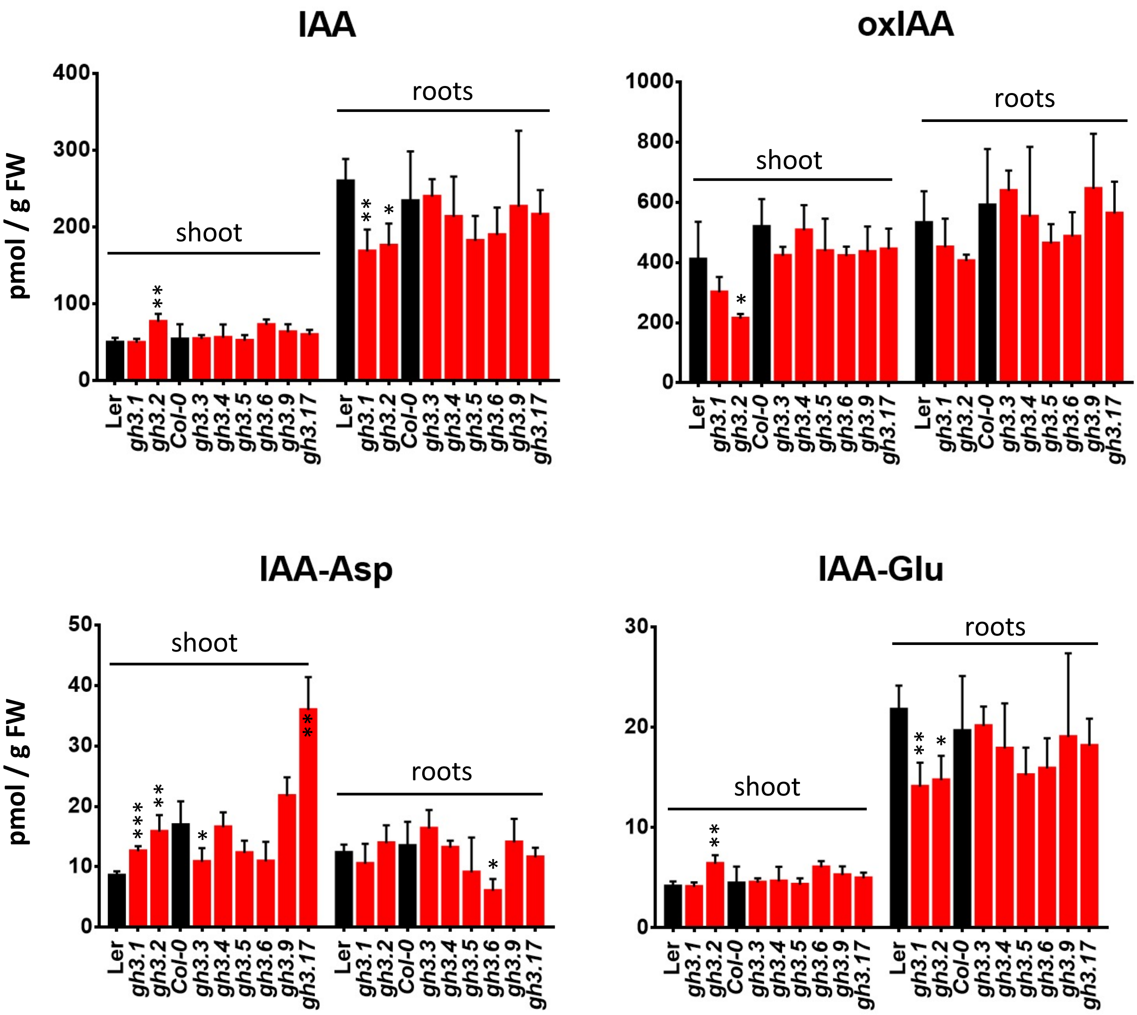


**Figure S7. Levels of IAA and IAA metabolites in shoots and roots from group-II *gh3* single mutants.** Concentration of indole-3-acetic acid (IAA), 2-oxindole-3-acetic acid (oxIAA), indole-3-acetyl-aspartate (IAA-Asp) and indole-3-acetyl-glutamate (IAA-Glu) in rosettes and roots from 7-day-old *gh3* single mutants (red bars) and their correspondent wild-type background (black bars). Error bars indicate standard deviation. Asterisks indicate significantly different concentration values compared to the corresponding wild type in a two-tailed Student´s *t*-test (*$p<0.05$; **$p<0.01$; $n = 4$). FW: fresh weight.

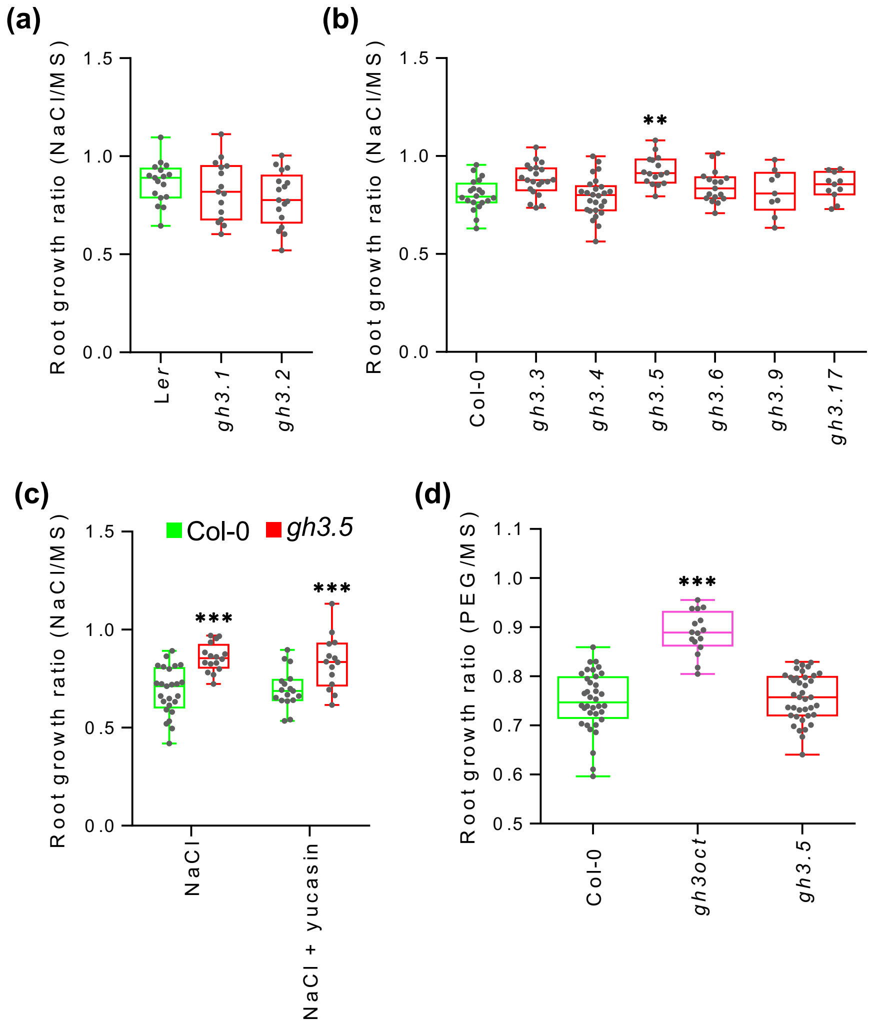


**Figure S8. Salinity and drought tolerance of *gh3* single mutants.** (a,b) Root growth ratio of *gh3* single mutants in (a) L*er* and (b) Col-0 background grown in 50 mM NaCl. To calculate the root growth ratio, the root length from seedlings on NaCl plates was divided by the mean root length of seedlings on MS control plates. (c) Effects of the YUCCA inhibitor yucasin on the *gh3.5* salt-tolerant phenotype. Seedlings were normally grown for 4 days and then transferred for another 3 days to media plates containing 50 mM NaCl or 50 mM NaCl + 20 µM of the YUCCA inhibitor yucasin. To calculate the ratio, the length of the root from seedlings on NaCl and NaCl + yucasin plates was divided by the mean root length of seedlings grown on control MS and MS + yucasin plates, respectively. (d) Root growth ratio of Col-0, *gh3oct* and *gh3.5* grown on media plates perfused with a 250 mg/L solution of PEG8000. 4-day-old seedlings were transferred to PEG-perfused or MS-perfused (control) plates and grown for another 2 days. To calculate the ratio, the root length from seedlings on PEG-containing plates was divided by the mean root length of seedlings on control plates. Asterisks indicate values significantly different from those of Col-0 in a two-tailed Student´s *t*-test (***p*<0.01, ****p*<0.001, *n* ≥ 14).

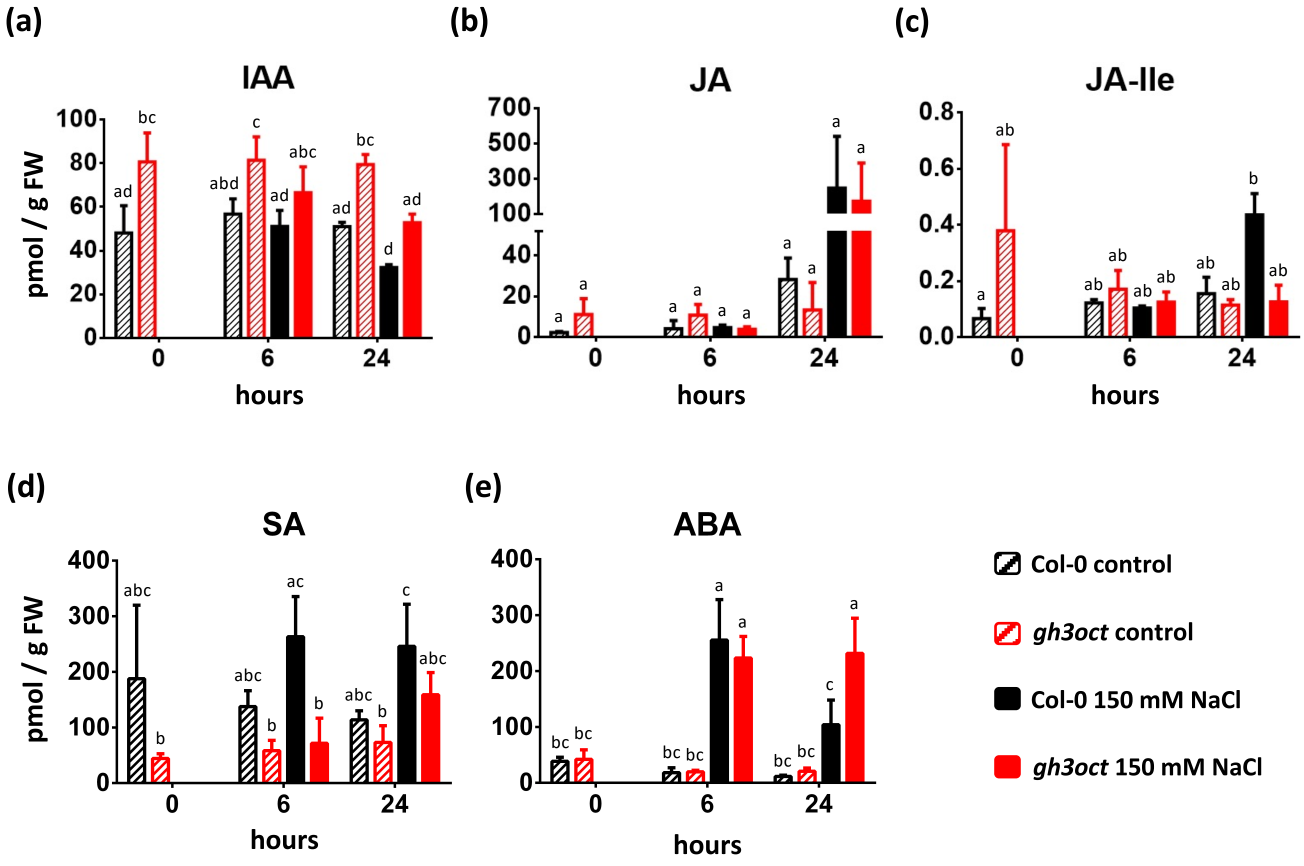


**Figure S9. Levels of IAA and stress-related phytohormones in response to salinity in the *gh3oct* mutant.** Concentrations of indole-3-acetic acid (IAA), jasmonic acid (JA), jasmonic acid-isoleucine (JA-Ile), salicylic acid (SA) and abscisic acid (ABA) in 9-day-old Col-0 (black bars) and *gh3oct* mutant (red bars) seedlings were determined 0, 6 and 24 hours after transferring to control MS plates (striped bars) or plates supplemented with 150 mM NaCl (solid bars). Error bars indicate the standard deviation. Different letters indicate significant differences ($p$<0.05) according to Tukey´s post-hoc test. FW: fresh weight.

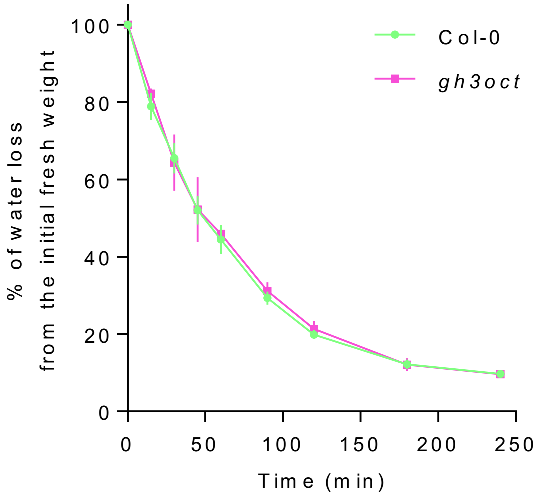


**Figure S10. Col-0 and *gh3oct* plants lose water at the same rate.** Aerial parts from 9-day-old Col-0 and *gh3oct* seedlings were detached and immediately weighed ($t$ = 0 min). The weight of the samples was thereafter inspected at the indicated times. Data points represent the mean ± standard deviation ($n$ = 4). No significant differences were found across the timepoints.

**Table S1**. Insertion lines combined to obtain the *gh3oct* mutant

| Gene | AGI code | Insertion | Background |
|---|---|---|---|
| *GH3.1* | AT2G14960 | SGT4174-5-2 | L*er* |
| *GH3.2* | AT4G37390 | SGT5949-5-3 | L*er* |
| *GH3.3* | AT2G23170 | SM.37350 | Col-0 |
| *GH3.4* | AT1G59500 | SALK_102549 | Col-0 |
| *GH3.5* | AT4G27260 | SALK_151766 | Col-0 |
| *GH3.6* | AT5G54510 | SALK_013458 | Col-0 |
| *GH3.9* | AT2G47750 | SM.23766 | Col-0 |
| *GH3.17* | AT1G28130 | SALK_050597 | Col-0 |

**Table S2**. Primer sets used in this work

| Purpose | Primer name | Primer sequence (5’ → 3’) | |
|---|---|---|---|
| | | Forward primer (F) | Reverse primer (R) |
| Genotyping | GH3.1 | GGATCCATGGCGGTAGACTCCAAC | AGGACAGAAAGGCGGTCAAG |
| | GH3.2 | GCTCCGACTCGTCCCAAAG | GTCGACCTAACGACGTCGTTCTGG |
| | Ds5'-1 | ACGGTCGGGAAACTAGCTCTAC | |
| | GH3.3 | CTATCGGTGACAGGCAGAGTC | AACCTCAGCATTTAGTCTTCACG |
| | GH3.9 | ATGGGTTTCGCACTTAGGAAG | TCAACGACTTTATCACAGGGG |
| | 3’-dSpm | TACGAATAAGAGCGTCCATTTTAGAGTGA | |
| | GH3.4 | CAATGACGGGATTTTGATCAC | TGTGGAGCGGAATTATGAAAC |
| | GH3.5 | AGGCCAGTGTTGTTGTCTTTG | TGGTCTTGAGCATAGATTCCG |
| | GH3.6 | AAACCTAAACGATGCCTGAGG | CTCAGGCCAATGTTTCTCAAG |
| | GH3.17 | GAAAAGTGAAAACTGTTGACGAG | CCTCTTGCTCACGGAGTACAC |
| | LBb1.3 | ATTTTGCCGATTTCGGAAC | |
| RT-qPCR | qGH3.1 | GGACAACTCGGTTGGACCAT | TGCACCTCTTGAGATTGCGT |
| | qGH3.2 | TGCGTGAGCTTCACACCTAT | CTAAAACCGCACATCATCCG |
| | qGH3.3 | CTCCGTGCCATTGGATTCCT | ATCAGCCAGTTCTTGGTCCG |
| | qGH3.4 | TGATCGGTGTGAGGCTTACG | TGGAGATACGTGTGGTGCAG |
| | qGH3.5 | TGCTCCAATTATCGAGCTATTGA | TGTTTGTGACCAGGAACCCA |
| | qGH3.6 | ACCTATGCTGGGCTTTACAGG | GCGGCATATGAAGCTGAACTG |
| | qGH3.9 | CGACGATGAACAAGTCCCCT | GGTAACGGTACAATCCTGCGA |
| | qGH3.17 | CATTTGTTAAGTTGCTAATTGGTGT | AGAGGACTTTGCTGAAAGTTTGT |
| | qACT2 | CCGCTCTTTCTTTCCAAGC | CCGGTACCATTGTCACACAC |
